# Relationship of maternal cytomegalovirus-specific antibody responses and viral load to vertical transmission risk following primary maternal infection in a rhesus macaque model

**DOI:** 10.1101/2023.04.21.537769

**Authors:** Claire E. Otero, Richard Barfield, Elizabeth Scheef, Cody S. Nelson, Nicole Rodgers, Hsuan-Yuan Wang, Matilda J. Moström, Tabitha D Manuel, Julian Sass, Kimberli Schmidt, Husam Taher, Courtney Papen, Lesli Sprehe, Savannah Kendall, Angel Davalos, Peter A. Barry, Klaus Früh, Justin Pollara, Daniel Malouli, Cliburn Chan, Amitinder Kaur, Sallie R. Permar

**Affiliations:** Department of Pathology, Duke University, Durham, NC; Department of Pediatrics, Weill Cornell Medical College, New York, NY; Department of Biostatistics and Bioinformatics, Duke University, Durham, NC; Tulane National Primate Research Center, Covington, LA; Division of Allergy and Clinical Immunology, Department of Medicine, Brigham and Women’s Hospital, Boston, MA; Duke Human Vaccine Institute & Department of Surgery, Duke University, Durham, NC; Department of Immunology, Duke University, Durham, NC; Department of Mathematics, North Carolina State University, Raleigh, NC; Center for Immunology and Infectious Diseases, University of California, Davis, CA; Vaccine and Gene Therapy Institute, Oregon Health & Science University, Beaverton, OR

## Abstract

Cytomegalovirus (CMV) is the most common congenital infection and cause of birth defects worldwide. Primary CMV infection during pregnancy leads to a higher frequency of congenital CMV (cCMV) than maternal re-infection, suggesting that maternal immunity confers partial protection. However, poorly understood immune correlates of protection against placental transmission contributes to the current lack of an approved vaccine to prevent cCMV. In this study, we characterized the kinetics of maternal plasma rhesus CMV (RhCMV) viral load (VL) and RhCMV-specific antibody binding and functional responses in a group of 12 immunocompetent dams with acute, primary RhCMV infection. We defined cCMV transmission as RhCMV detection in amniotic fluid (AF) by qPCR. We then leveraged a large group of past and current primary RhCMV infection studies in late-first/early-second trimester RhCMV-seronegative rhesus macaque dams, including immunocompetent (n=15), CD4+ T cell-depleted with (n=6) and without (n=6) RhCMV-specific polyclonal IgG infusion before infection to evaluate differences between RhCMV AF-positive and AF-negative dams. During the first 3 weeks after infection, the magnitude of RhCMV VL in maternal plasma was higher in AF-positive dams in the combined cohort, while RhCMV glycoprotein B (gB)- and pentamer-specific binding IgG responses were lower magnitude compared to AF-negative dams. However, these observed differences were driven by the CD4+ T cell-depleted dams, as there were no differences in plasma VL or antibody responses between immunocompetent AF-positive vs AF-negative dams. Overall, these results suggest that levels of neither maternal plasma viremia nor humoral responses are associated with cCMV following primary maternal infection in healthy individuals. We speculate that other factors related to innate immunity are more important in this context as antibody responses to acute infection likely develop too late to influence vertical transmission. Yet, pre-existing CMV glycoprotein-specific and neutralizing IgG may provide protection against cCMV following primary maternal CMV infection even in high-risk, immunocompromised settings.

**Author summary:** Cytomegalovirus (CMV) is the most common infectious cause of birth defects globally, but we still do not have licensed medical interventions to prevent vertical transmission of CMV. We utilized a non-human primate model of primary CMV infection during pregnancy to study virological and humoral factors that influence congenital infection. Unexpectedly, we found that the levels virus in maternal plasma were not predictive of virus transmission to the amniotic fluid (AF) in immunocompetent dams. In contrast, CD4+ T cell depleted pregnant rhesus macaques with virus detected in AF had higher plasma viral loads than dams not showing placental transmission. Virus-specific antibody binding, neutralizing, and Fc-mediated antibody effector antibody responses were not different in immunocompetent animals with and without virus detectable in AF, but passively infused neutralizing antibodies and antibodies binding to key glycoproteins were higher in CD4+ T cell-depleted dams who did not transmit the virus compared to those that did. Our data suggests that the natural development of virus-specific antibody responses is too slow to prevent congenital transmission following maternal infection, highlighting the need for the development of vaccines that confer levels of pre-existing immunity to CMV-naïve mothers that can prevent congenital transmission to their infants during pregnancy.

## Introduction

Cytomegalovirus (CMV) is the most common congenital infection and the leading non-genetic cause of hearing loss in pediatric populations (1, 2). Placental transmission following primary human CMV (HCMV) infection is much more common (30–40%) than after non-primary infection (3–4%), suggesting that the maternal immune system is effective in reducing the risk of vertical CMV transmission (3). Furthermore, the biological factors contributing to or determining the likelihood of experiencing a congenital infection to the fetus in a subpopulation of CMV naïve women who contract a primary infection during pregnancy without having pre-existing CMV specific immunity remain poorly defined. Identification of immune responses that can naturally protect against congenital CMV (cCMV) would be enormously valuable for the rational design of an effective vaccine to prevent vertical CMV transmission.

The objective of this study was to identify maternal humoral immune responses associated with vertical CMV transmission in the high-risk setting of primary maternal infection, as a strategy to guide vaccine development. The study of primary CMV infection faces several logistical hurdles. First, primary CMV infection is difficult to detect in humans because CMV infection is typically asymptomatic or exceedingly mild in healthy individuals, including during pregnancy (4). This diagnostic dilemma makes human studies of early immune responses to infection challenging. Unfortunately, HCMV cannot be used to infect any animal model commonly used in the laboratory to study virus pathogenesis as all examined CMV species exhibit a high degree of species-specificity and can hence only productively replicate in the host species they have co-evolved with (5, 6). Moreover, not all animal CMVs cross the placenta, which may be a result of differences in placental biology or viral characteristics, further limiting animal models for the study of cCMV and placental CMV transmission(7, 8). Non-human primates (NHPs) bear the closest resemblance to humans in terms of anatomy and physiology of pregnancy, and immune responses to viral infection (9, 10, 11, 12). Furthermore, as CMV evolution has been shown to track host evolution(13), NHP CMVs represent the closest relatives to Great Ape and human CMVs as they share the most recent common ancestor, and conservation of coding ORFs and gene families, viral immune evasion strategies and viral pathogenesis has been demonstrated (14). Thus, we have developed a rhesus macaque model of primary, maternal CMV infection during the early stages of pregnancy, at the intersection of the first and second trimesters of gestation. In this model, congenital transmission occurs a fraction of the time in immunocompetent dams and universally in CD4+ T cell depleted dams without additional intervention (15), which has proven useful in defining the immunological and virological determinants of congenital CMV transmission (16).

We hypothesized that high magnitude maternal viremia is associated with vertical RhCMV transmission, particularly in this setting of primary maternal infection where no pre-existing immunity exists. We reasoned that increased virus circulation in the maternal blood should result in a greater viral load at the maternal-fetal interface, increasing the likelihood of placental infection and congenital transmission. This idea has recently been supported by mathematical modeling of vertical CMV transmission (17). Furthermore, we hypothesized that maternal virus-specific functional antibody responses such as neutralization, antibody dependent cellular cytotoxicity (ADCC), and antibody dependent cellular phagocytosis (ADCP) are important for controlling viral replication and preventing vertical transmission, as cell-to-cell spread is a key mechanism for CMV dissemination (18), meaning that antibody functions that target both cell-free and cell-associated virus may be needed for adequate control.

We leveraged multiple RhCMV-seronegative, pregnant NHP groups that were challenged with RhCMV at the end of the first/early second trimester of pregnancy from prior and current studies modeling vertical CMV transmission: a cohort of 15 total immunocompetent dams (3 dams from a historical study and 12 dams from a current study), 6 dams that received a single dose of CD4 T cell-depleting antibody 1 week prior to infection, and 6 total dams that were CD4+ T cell depleted (CD4-depleted) 1 week prior to infection and received a polyclonal RhCMV-specific IgG infusion just prior to infection to model pre-existing antibody (15, 16). We began by characterizing the antibody responses in a novel cohort of 12 immunocompetent, acutely RhCMV-infected dams, including ADCP and ADCC functions, which had not been previously measured in the context of RhCMV infection. Many of these responses have been previously reported for the historical cohorts, and available plasma samples from these studies were re-analyzed for recently developed functional antibody assays. Subsequent analysis of these responses and maternal viral load (VL) revealed no significant differences in the magnitude of these variables in immunocompetent dams that did or did not have detectable virus in amniotic fluid (AF), but RhCMV glycoprotein-specific binding responses and neutralizing antibody responses were higher magnitude in CD4-depleted animals that received passive IgG infusion and did not have viral DNA detectable in AF compared to those that did.

## Results

### RhCMV vertical transmission confirmed in 5/12 novel immunocompetent dams following primary RhCMV infection during pregnancy by detection of virus in AF

We challenged 12 RhCMV-seronegative rhesus dams via intravenous injection of a combination of 1.0×10^6^ PFU of UCD52 strain and 1.0×10^6^ PFU of a bacterial artificial chromosome (BAC)-derived, clonal full-length (FL) RhCMV (13). We monitored VL of maternal plasma and AF following RhCMV infection using qPCR detection of viral DNA targeting the RhCMV IE gene(19). Plasma VL peaked 1–2 weeks post infection in all dams, and many had detectable VL through 14 weeks post infection (**Figure 1B**). Five of the twelve dams in this cohort had CMV DNA detectable by qPCR in the AF collected weekly between 2–12 weeks after infection, confirming vertical transmission (AF-positive). AF virus was detected within 4 weeks post infection in three dams and after 10 weeks post infection in the remaining two AF-positive dams (**Figure 1C**). Importantly, the kinetics of the plasma VL was similar in AF-positive vs AF-negative dams.

**Figure 1:**
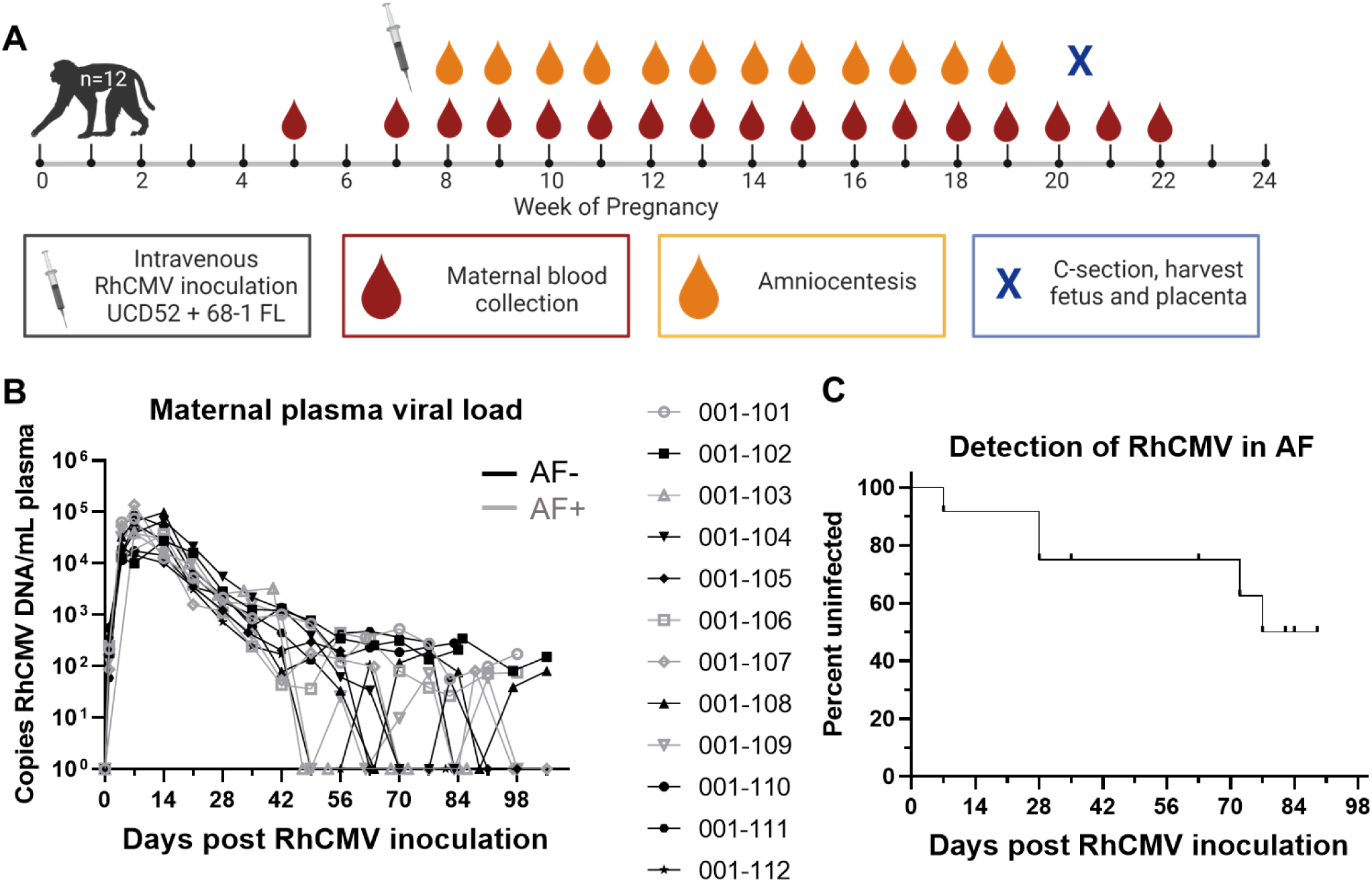
Plasma VL and vertical transmission events following primary maternal RhCMV infection in immunocompetent dams (n = 12) (A) The study schedule for a new cohort of 12 RhCMV seronegative, pregnant rhesus macaques receiving intravenous inoculation of 1×10^6^ pfu of two RhCMV strains (clinical isolate UCD52 and cloned full length, FL, 68.1 viral variants) in late first/early second trimester, followed by weekly blood draws and amniocentesis until hysterotomy near term at 20–22 weeks gestation, prior to fetal viability. A pre-infection blood draw was collected between 1–4 weeks before infection. (B) Maternal plasma VL kinetics measured by qPCR against the IE-1 gene for RhCMV-positive amniotic fluid (AF-positive, n = 5) and AF-negative (n = 7) dams. Data shown as the mean value of 6 or more technical replicates for each sample. (C) Survival curve showing the timing of detection of vertical transmission, defined as amniotic fluid RhCMV qPCR result above the limit of detection in 2 or more of 12 technical replicates. Vertical transmission was confirmed in 5/12 dams by this definition.

### RhCMV-specific antibody binding kinetics following primary RhCMV infection during pregnancy

Plasma RhCMV-specific IgM responses were detectable at low levels by 2 weeks post infection and remained elevated throughout the pregnancy (**Figure 2A**). IgG binding to whole RhCMV virions and key envelope glycoproteins, glycoprotein B (gB) and the pentameric complex (PC), shared similar kinetics, with detectable responses by 2 weeks post infection in most animals, with a sharp increase in antibody titers between 2–4 weeks post infection followed by a plateau or a more gradual increase in antibody levels throughout the remainder of the pregnancy (**Figure 2B–E**). Notably, we measured gB-specific IgG binding to both soluble ectodomain via ELISA and cell-associated gB via a transfected cell binding assay, and each demonstrated similar kinetics (**Figure 2C, D**). Overall, these responses are largely consistent with the expected binding antibody responses during the acute phase of infection in humans (20).

**Figure 2:**
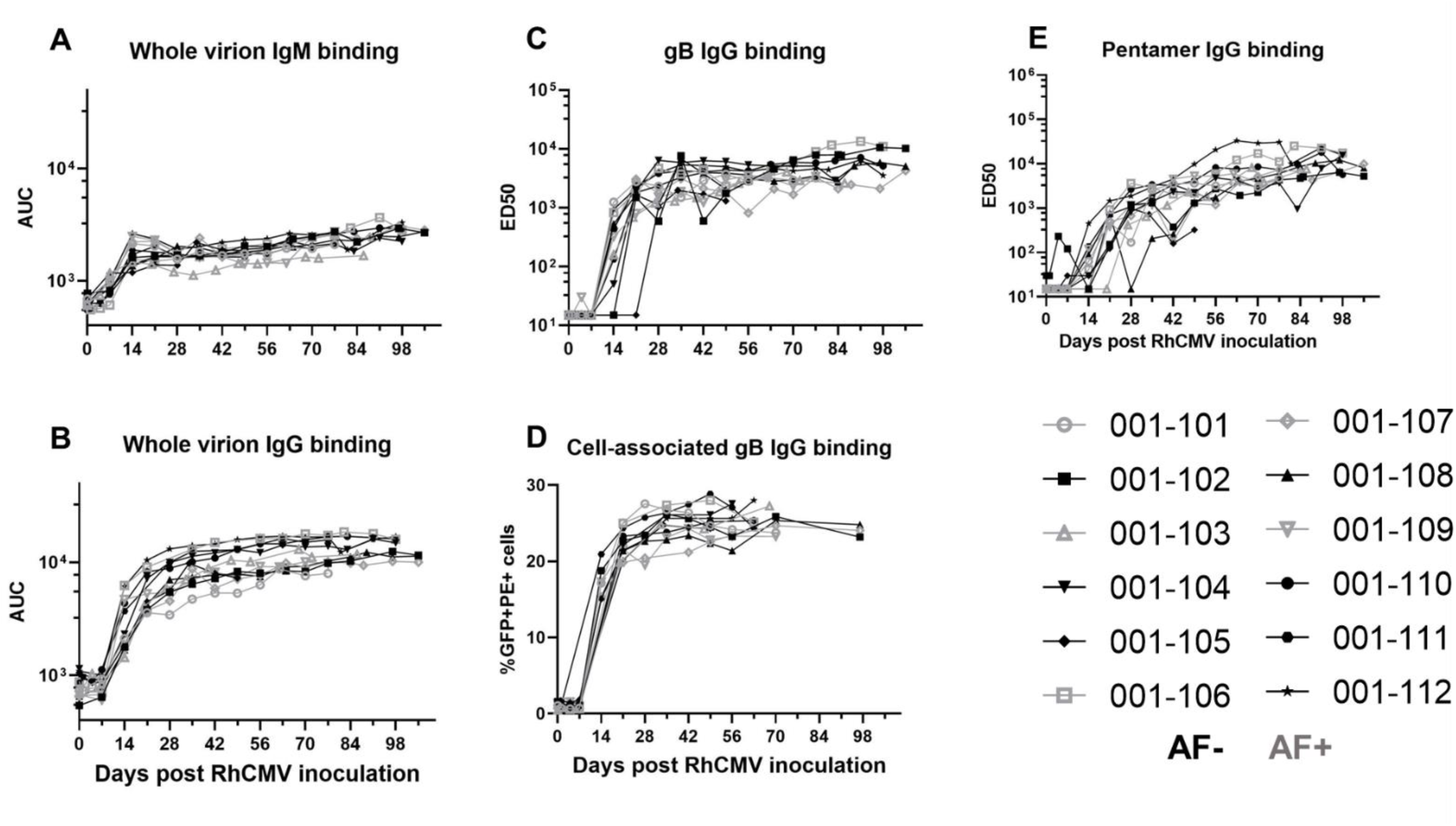
Plasma RhCMV-specific IgM and IgG binding responses following acute RhCMV infection during pregnancy. In a novel cohort of 12 RhCMV seronegative, immunocompetent dams primarily infected with RhCMV at the end of first/early second trimester of pregnancy, we measured the following RhCMV-specific antibody binding responses longitudinally: (A) IgM binding to whole UCD52 RhCMV virions, (B) IgG binding to whole UCD52 virions, (C) IgG binding to the ectodomain of UCD52 gB, (D) IgG binding to cell-associated UCD52 gB, and (E) IgG binding to soluble PC. AF-positive dams are shown in grey, and AF-negative dams are shown in black. Data shown as the mean of technical duplicates.

### RhCMV-specific antibody effector function kinetics following primary RhCMV infection during pregnancy

Similar to the kinetics of the RhCMV-specific IgG binding responses, RhCMV-neutralizing responses in both fibroblast and epithelial cells developed within 3 weeks post infection, reached an inflection point by 6 weeks post infection, and then gradually increased over the remainder of pregnancy (**Figure 2A, B**). In contrast, non-neutralizing Fc-mediated antibody effector responses, ADCP (antibody-mediated uptake by THP-1 monocytes) and ADCC (measured as antibody-mediated NK cell degranulation)(21), generally developed more gradually over the course of the infection and never reach a true peak during the study period (**Figure 3C, D**). Importantly, the kinetics of all antibody responses were similar in AF-positive and AF-negative dams (**Figures 2** and **3**).

**Figure 3:**
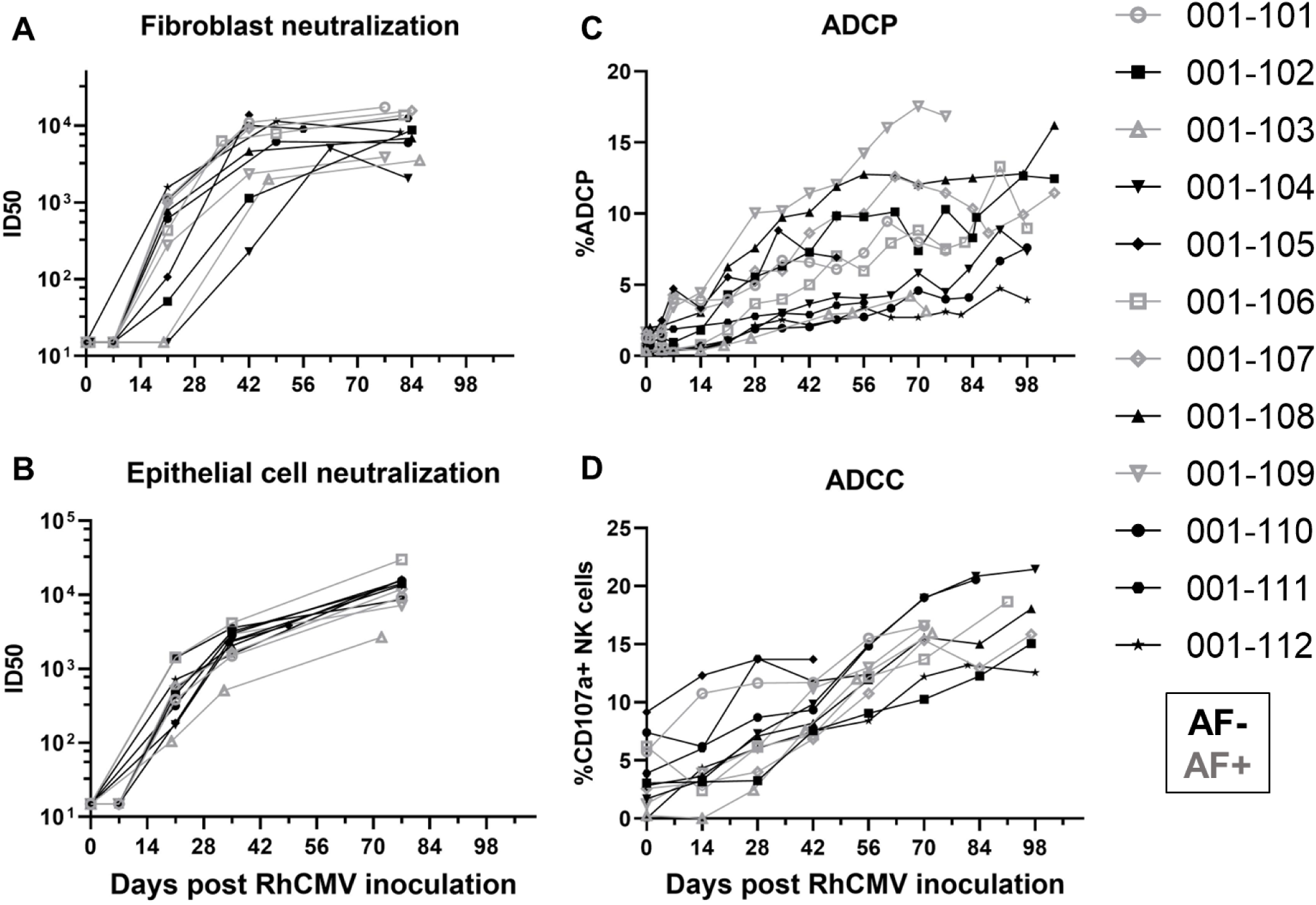
The kinetics of functional RhCMV-specific antibody responses in plasma during acute maternal infection. We measured functional antibody responses longitudinally in a novel cohort of 12 immunocompetent dams with primary RhCMV infection during pregnancy. (A) Neutralization of FL RhCMV on telomerized rhesus fibroblasts (TeloRFs), (B) neutralization of UCD52 RhCMV on monkey kidney epithelial cells, (C) monocyte-mediated antibody dependent cellular phagocytosis against free UCD52 RhCMV, (D) NK cell degranulation, a component of cell-mediated antibody dependent cellular cytotoxicity against UCD52-infected TeloRFs. Data shown as the mean of technical duplicates.

### Inclusion of historical CD4-depleted and IgG-infused groups introduces perturbations of measured responses

For further analysis, we incorporated data from past studies of rhesus dams with primary RhCMV infection in late first / early second trimester (15, 16). The timing of maternal infection, administration of RhCMV, and total inoculum dose are similar among all of these groups, but the virus strains and immunologic manipulations differ somewhat among the groups (**Figure 4**). As reported in these previous studies, CD4-depletion results in higher maternal plasma viral load, consistent vertical RhCMV transmission, and a high rate of fetal loss compared to immunocompetent dams (15). However, pre-existing, potent antibody infusion leads to a reduction in maternal plasma viral load and demonstrated protection in this extremely high risk setting of CD4-depletion (16). We chose to include animals from these previous studies to bolster the total sample size and introduce the following perturbations to the measured responses: 1) maternal plasma viral load was higher in CD4-depleted animals and lower in CD4-depleted animals receiving a potently-RhCMV neutralizing and dose-optimized IgG infusion, 2) humoral responses were generally delayed in CD4-depleted dams, and 3) dams receiving passive IgG infusion before infection had high levels of RhCMV-specific IgG in the first week (**Figure 5**). With the understanding that treatment group is a potential confounding factor, we performed subgroup analyses for our primary analysis of pairwise comparisons between AF-positive and AF-negative dams.

**Figure 4:**
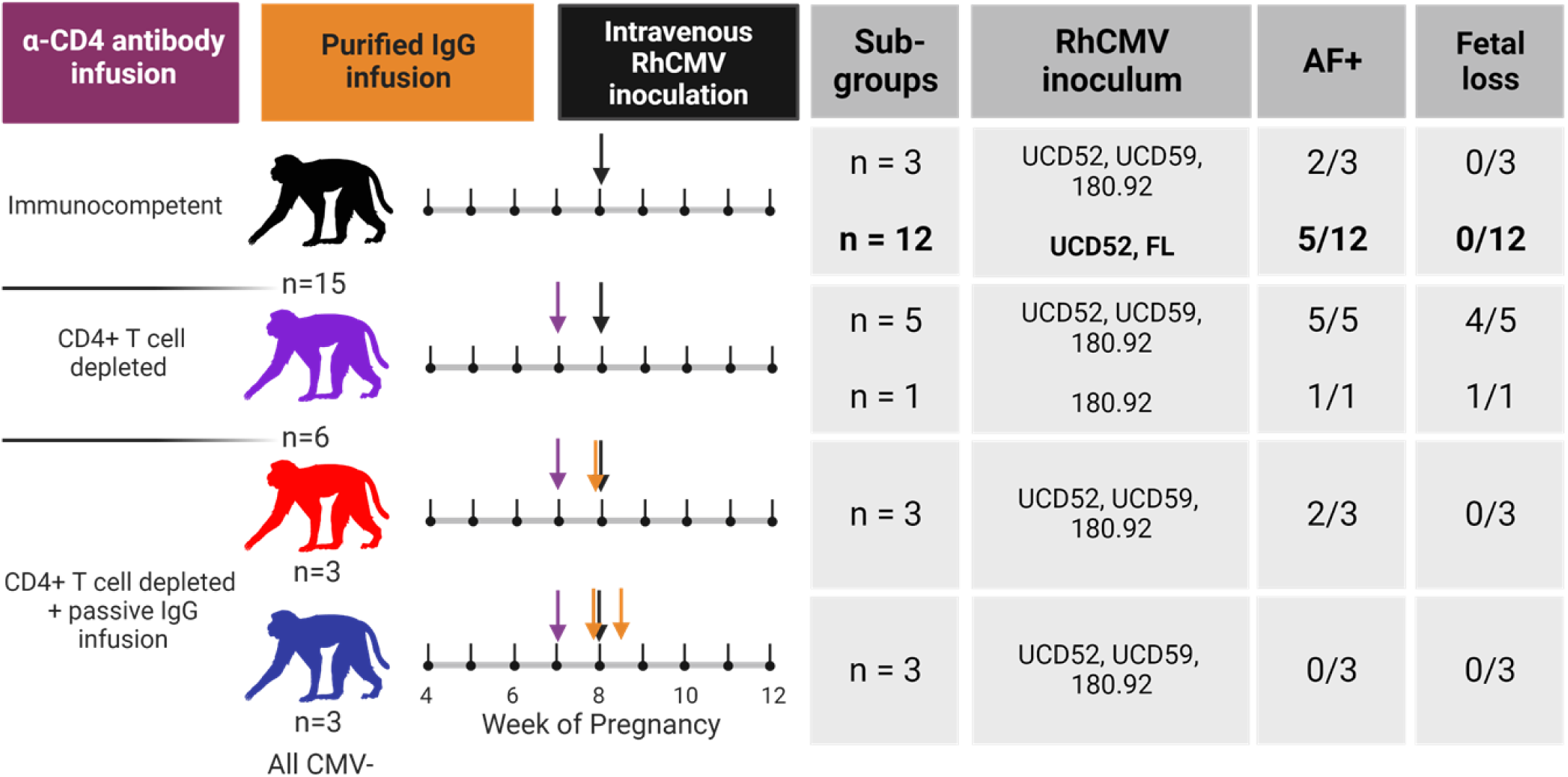
Combination of RhCMV seronegative, pregnant rhesus macaque cohorts. Treatment schedules for each of the rhesus macaque cohorts included in the combined analysis. A new group of 12 immunocompetent dams is highlighted in bold, and all other groups are from historical studies completed between 2013–2016 (15, 16). All dams were infected by intravenous inoculation of 1×10^6^ pfu for each UCD52, UCD59, and FL RhCMV and 2–3×10^6^ TCID_50_ for 180.92 at week 7–9 of gestation. A total of 15 immunocompetent dams (black) included 3 dams from a previous study in which animals received a 1:1:2 combination of low passage isolates UCD52 and UCD59 and lab adapted strain 180.92 as well as 12 dams inoculated with a 1:1 combination of UCD52 and BAC-derived full length (FL) RhCMV. A total of 6 dams received a 50 mg/kg dose of recombinant rhesus anti-CD4+ T-cell–depleting antibody 1 week prior to infection. Five of these dams were infected with 1:1:2 UCD52, UCD59, and 180.92, and the remaining dam was infected with 180.92 alone. Six additional animals received CD4-depleting antibody 1 week before infection and passive infusion of a polyclonal RhCMV-positive IgG preparation before infection with 1:1:2 UCD52, UCD59, and 180.92. IgG for passive infusion was purified from RhCMV-seropositive plasma donors screened for high RhCMV binding (red) or high RhCMV neutralization on epithelial cells (blue). The high RhCMV binding IgG infusion was given at a single 100 mg/kg dose 1 hour before infection, and the high RhCMV neutralizing IgG infusion was given in a two-dose, optimized regimen with the initial 150 mg/kg dose 1 hour prior to infection and the second 100 mg/kg dose 3 days post infection. Of the immunocompetent dams, 7/15 had detectable RhCMV DNA in amniotic fluid. All 6 of the CD4-depleted dams were AF-positive, and 5 experienced spontaneous abortion. Two of the dams receiving the high RhCMV binding IgG infusion and none of the dams receiving the high RhCMV neutralizing IgG infusion were AF-positive.

**Figure 5:**
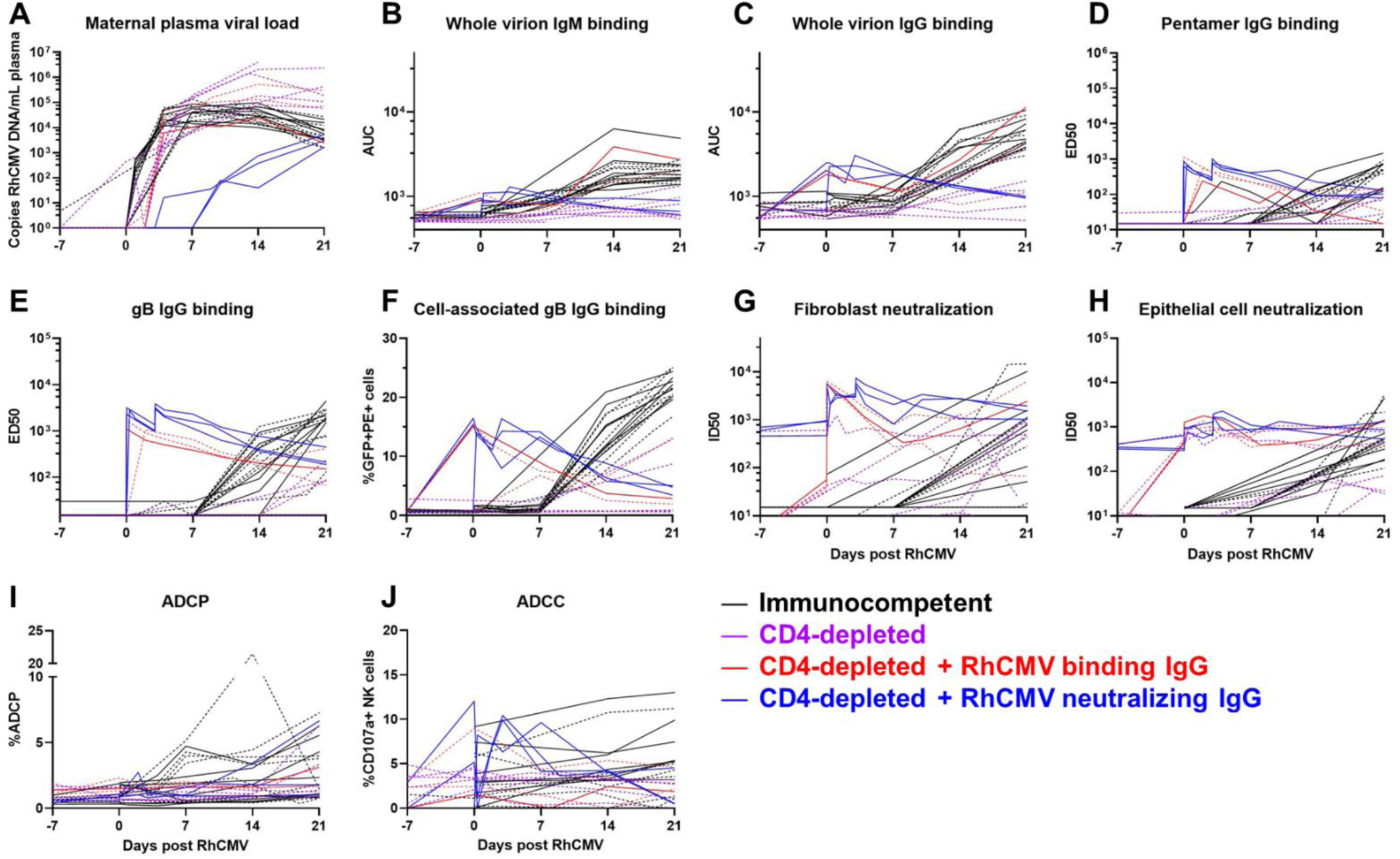
Kinetics of plasma VL and RhCMV-specific antibody responses for all treatment groups through 3 weeks post RhCMV infection. (A) Maternal plasma VL measured by qPCR for the IE-1 gene; (B) IgM and (C) IgG binding to whole UCD52 RhCMV virions; IgG binding to (D) PC, (E) soluble gB ectodomain by ELISA, and (F) cell-associated gB via transfected cell binding assay; antibody mediated RhCMV neutralization on (G) fibroblasts and (H) epithelial cells; (I) ADCP; and (J) ADCC. Treatment groups are represented by the color of each line. Solid lines represent AF-negative dams, and dashed lines represent AF-positive dams. Previously published data for historical studies was used for soluble gB- and PC-binding IgG responses and fibroblast and epithelial cell neutralization (15, 16), and available samples from those studies were utilized for measurement of remaining assays.

### The magnitude of the plasma VL is not significantly different between AF-positive versus AF-negative immunocompetent dams

To distill each virological and immunological variable for comparison across vertically transmitting and non-transmitting dams, we calculated the area-under-the-curve (AUC) for the first three weeks post infection, which is the time period when vertical RhCMV transmission most often occurs and each measured response develops. We had no evidence to suggest that maternal VL was different between AF-positive vs AF-negative immunocompetent dams (Wilcoxon p_unadj_ = 0.69, p_FDR_ = 0.87). In stark contrast, plasma VL was higher magnitude in AF-positive dams in the CD4-depleted groups of dams, with AF-positive dams demonstrating higher maternal plasma VL (p_unadj_ = 0.004, p_FDR_ = 0.004), driving the difference in the maternal plasma VLs in the combined group (p_unadj_ = 0.0013, p_FDR_ = 0.0013). However, the differences in plasma VL in CD4-depleted dams is largely confounded by the treatment group, as all four AF-negative CD4-depleted animals received passive IgG infusion before infection and all six of the CD4-depleted dams without pre-existing antibody were AF-positive (**Figure 4**). Thus, pre-existing CMV humoral immunity may have contributed directly to the reduction in plasma VL and the lack of vertical CMV transmission.

### Humoral immune responses are not statistically significantly different between AF-positive and AF-negative immunocompetent dams, yet plasma cell-associated gB and PC-specific IgG responses were higher magnitude in nontransmitting dams from the entire cohort

As observed for maternal plasma VL, none of the RhCMV-specific antibody binding or functional antibody responses were statistically different in AF-positive versus AF-negative immunocompetent dams (**Table 1, Figure 6**). In comparing RhCMV-specific humoral responses across the entire dam cohort, the responses that were statistically associated with transmission status were plasma IgG binding to cell-associated gB (p_unadj_ = 0.017, p_FDR_ = 0.04) and soluble PC (p_unadj_ = 0.028, p_FDR_ = 0.0.047). Within the CD4-depleted groups, plasma RhCMV glycoprotein-specific binding IgG responses and neutralization in both fibroblasts and epithelial cells were statistically higher in AF-negative compared to AF-positive dams by unadjusted Wilcoxon Rank Sum tests (**Table 1**).

**Figure 6:**
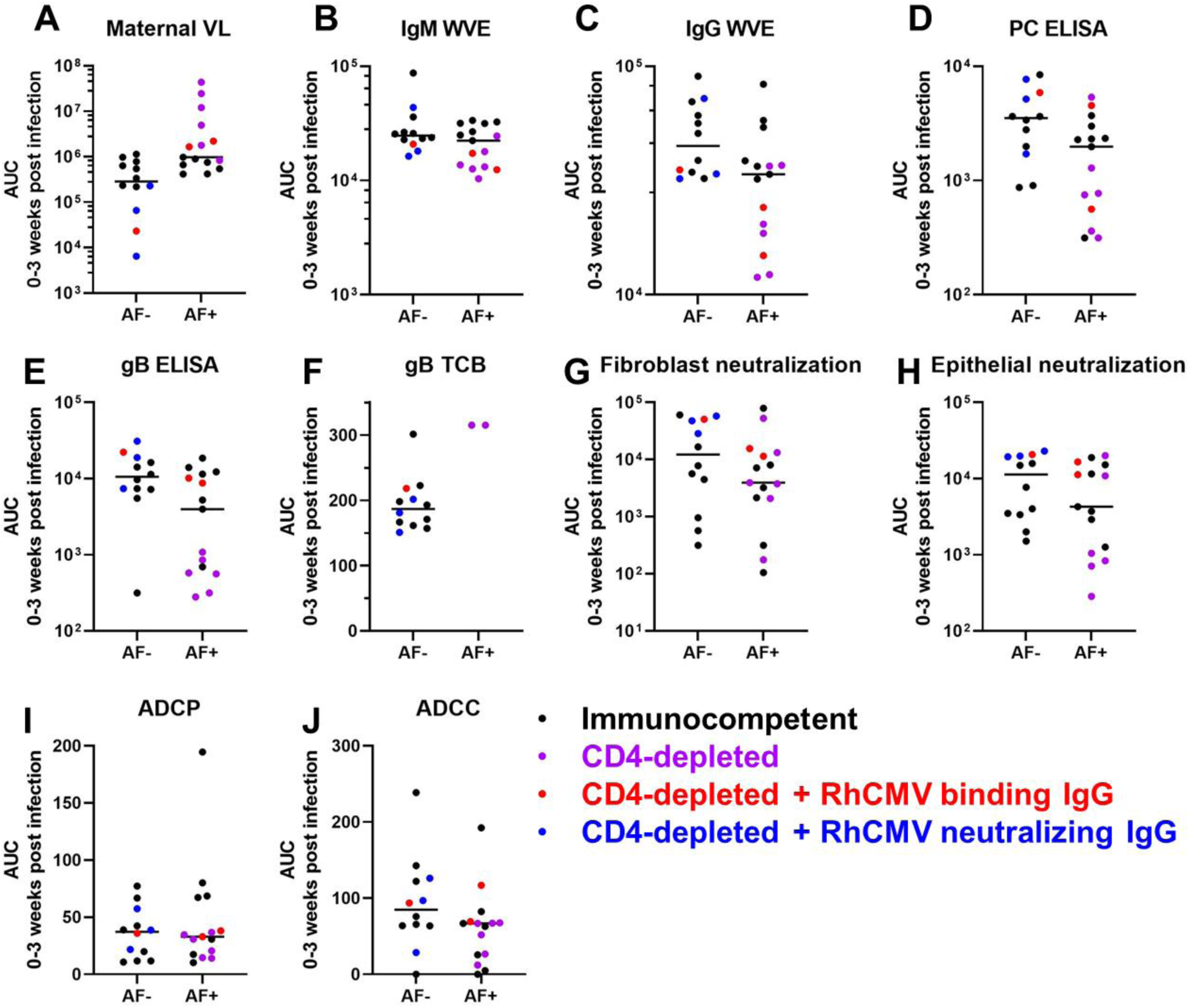
Univariate comparison of maternal VL and humoral responses by vertical transmission status. The area-under-the-curve of each response over 0–3 weeks post RhCMV infection was used to distill the magnitude and kinetics of each response into a single value in a critical window for vertical transmission. Amniotic fluid (AF) was assessed for RhCMV DNA by qPCR and a positive result (AF-positive) was defined by 2 or more of 12 replicates with detectable amplification between 2 and 12 weeks after maternal infection. Pairwise comparisons between AF-positive and AF-negative dams are shown for all treatment groups combined, with symbol color representing each treatment group. AF-positive and AF-negative dams were compared using Wilcoxon Rank Sum Tests for the entire cohort as well as sub-group analyses of immunocompetent and CD4-depleted groups (**Table 1**).

**Table 1:** Univariate analysis comparing AF-positive versus AF-negative dams within subgroups.

|  |  |  |  |
| --- | --- | --- | --- |
| **Response** | **$p_{\text{unadjusted}}$ ($p_{\text{FDR}}$)** |  |  |
|  | **Immunocompetent (n=15)** | **CD4-depleted (n=12)** | **Combined cohort (n=27)** |
| Maternal plasma viral load | 0.694 (0.868) | **0.00404 (0.0404)** | **0.00128 (0.0128)** |
| Cell-associated gB binding | 0.397 (0.868) | **0.0162 (0.0404)** | **0.0148 (0.074)** |
| PC binding (ELISA) | 0.536 (0.868) | **0.0283 (0.0471)** | **0.0407 (0.136)** |
| gB binding (ELISA) | 0.955 (0.955) | **0.0162 (0.0404)** | 0.0601 (0.149) |
| Whole virion IgG binding | 0.463 (0.868) | 0.109 (0.136) | 0.0743 (0.149) |
| Neutralization on epithelial cells | 0.694 (0.868) | **0.0162 (0.0404)** | 0.114 (0.168) |
| ADCC | 0.351 (0.868) | 0.214 (0.214) | 0.118 (0.168) |
| Neutralization on fibroblasts | 0.805 (0.894) | **0.0283 (0.0471)** | 0.135 (0.168) |
| Whole virion IgM binding | 0.613 (0.868) | 0.0727 (0.104) | 0.376 (0.418) |
| ADCP | 0.336 (0.868) | 0.154 (0.171) | 0.886 (0.886) |
| P-values show exact results from Wilcoxon Rank Sum Tests as unadjusted p-value (FDR p-value). **Bold** indicates P-value <0.05 and FDR p-value <0.2. Abbreviations: Amniotic fluid (AF), False Discovery Rate corrected (FDR); immunoglobulin M (IgM), immunoglobulin G (IgG), glycoprotein B (gB), enzyme-linked immunosorbent assay (ELISA), pentameric complex (PC), antibody dependent cellular phagocytosis (ADCP), antibody dependent cellular cytotoxicity (ADCC). |  |  |  |

### Plasma RhCMV-specific antibody responses are indirectly related to plasma VL

Using Spearman’s rank correlations, we found that all of the measured RhCMV-specific antibody responses demonstrated weak negative associations with maternal plasma VL across the entire cohort (**Figure 7A**). Further, we found moderately strong positive associations between IgG and IgM binding to whole UCD52 virions (r = 0.74, 95% CI: 0.44, 0.91), soluble gB and PC IgG binding (r = 0.83, 95% CI: 0.64, 0.91), and between fibroblast and epithelial cell neutralization responses (r = 0.62, 95% CI: 0.23, 0.86). There were also moderate positive associations between the glycoprotein-specific IgG binding and neutralization responses (**Figure 6**), which is unsurprising given that gB and PC are key targets of CMV-neutralizing antibodies (22, 23). Additionally, principal component analysis (PCA) suggesting a clustering of transmitters away from non-transmitters, largely driven by the loading for maternal plasma VL, which was negatively correlated with humoral responses (**Figure 7B**). However, there is a clear influence of group status as all of the CD4-depleted dams are clustering together (**Figure 7C**).

**Figure 7:**
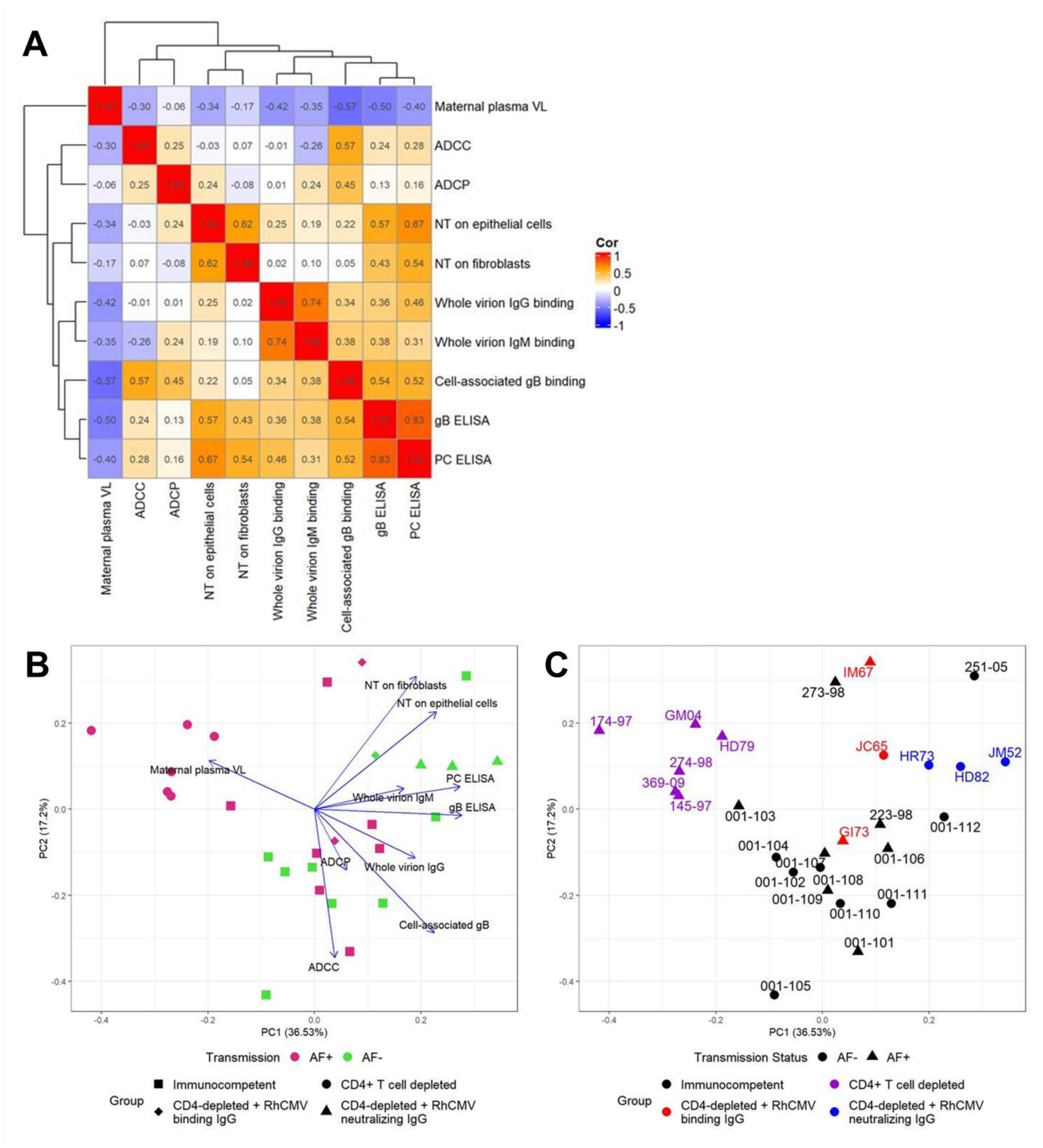
RhCMV-specific antibody responses negatively associate with maternal VL. (A) Correlation matrix showing Spearman correlation coefficients for pairwise correlations between the AUC of each RhCMV-specific antibody response over the first 3 weeks post infection. (B) Principal component analysis (PCA) using the AUC of each response over the first 3 weeks post infection, showing clustering by AF-negative (magenta) versus AF-positive (green) with treatment groups represented by the symbol shape. Loading vectors for each measured response demonstrate the influence of each response. (C) The same PCA with group represented by color, transmission status represented by shape, and animal IDs labeled to better demonstrate the effect of treatment group.

## Discussion

Despite the high risk setting of primary maternal CMV infection during pregnancy, only a minority (30-40%) of these cases will congenitally transmit the virus to their fetus. The determinants of cCMV transmission following acute maternal CMV infection remain largely unknown, yet could be an important guide to vaccine development. Human studies have previously suggested that high maternal anti-CMV ADCP responses is associated with protection from vertical HCMV transmission (24). Yet, human studies are inherently limited by the inability to accurately time primary CMV infection given its often mild or silent presentation. Thus, the rhesus macaque model of cCMV transmission following acute CMV infection in pregnancy represents a unique opportunity to define the early kinetics of virus replication and immunity, as well as their impact on vertical transmission risk.

Importantly, the rhesus model of cCMV transmission following primary RhCMV infection at the end of 1^st^ trimester in immunocompetent dams and utilizing the end point of virus detected in AF fluid between 2 and 12 weeks after infection accurately modeled the epidemiology of human CMV transmission following acute infection during pregnancy, with 5 of 12 dams (41.7%) with virus detection in AF (3). Yet, contrary to our expectations and challenging a common hypothesis in the field (17, 25), we found that maternal plasma VL magnitude and kinetics were not significantly different between AF-positive and AF-negative immunocompetent dams. While the exact mechanism of transplacental CMV transmission has not been fully defined, the notion that more virus circulating in maternal blood leads to greater risk of transmission due to the increased interaction of CMV with the maternal-fetal interface is not supported by these results. Yet, there was a notable difference in the plasma VL of vertically transmitting and non-transmitting dams in the setting of extreme immune suppression via CD4+ T cell depletion. Notably, the viral loads of the immunocompetent animals tended to fall in the middle of the overall distribution, and we speculate that the level of maternal viremia may not be a useful predictor of vertical transmission based on this result. However, controlling maternal viremia to a lower level as we saw in some animals treated with a polyclonal CMV-targeting IgG infusion, is likely still a worthwhile goal towards the prevention of cCMV. Furthermore, we inoculated these monkeys via intravenous injection with the express purpose of modeling maternal viremia, but oral inoculation, which better models natural routes of infection may yield greater variation in viral load in immunocompetent animals (26). Our findings may provide insight as to why randomized trials of CMV hyperimmune globulin did not demonstrate reduced congenital transmission and may indicate that post-exposure prophylaxis strategies will have limited efficacy due to a small window of opportunity to prevent vertical transmission in the setting of primary maternal infection (27, 28). Yet, this finding is likely limited only to infection in the absence of pre-existing immunity and does not diminish the importance of vaccine-elicitation of pre-existing immunity that will reduce the risk of maternal infection and/or congenital transmission.

Also contrary to our expectations, we found no statistically significant differences in humoral immune responses in immunocompetent dams who did and did not have virus detected in AF. We speculate that because transmission often occurs shortly following infection in this setting of primary maternal infection, and the humoral response develops too slowly to protect from vertical transmission. Other characteristics of the maternal immune landscape, potentially related to innate immunity or T cell responses, may play more of a role in determining vertical CMV transmission in the context of acute, primary infection. However, as demonstrated by the CD4-depleted dams, pre-existing virus-specific IgG antibodies can protect from vertical RhCMV transmission (16), suggesting that immunization or pre-exposure prophylactic immunotherapeutic strategies are likely to be the most protective approach.

In combining the primarily-infected rhesus macaque dams for enhanced variability of antibody levels and power to detect differences in transmitting and non-transmitting dams, only maternal plasma RhCMV gB and PC glycoprotein-specific binding IgG responses were higher magnitude in those dams that did not have virus detected in AF compared to those that did. Interestingly, the cell-associated gB-binding IgG response was statistically different between the groups, but not against soluble gB, paralleling our previous finding that this response was an immune correlate of protection against virus acquisition in gB/MF59 vaccine studies (29). However, these differences are largely driven by the CD4-depleted groups, as similar trends are observed when we look exclusively at the CD4-depleted groups but are not present in the immunocompetent group. Neutralizing antibodies also stand out higher magnitude in non-transmitting compared to transmitting dams when focused on the CD4-depleted groups, which is unsurprising given the correlations between gB- and PC-binding IgG and neutralization responses. Furthermore, in the CD4-depleted group, dams receiving potent RhCMV-neutralizing IgG infusion produced from RhCMV-seropositive plasma donors screened for high epithelial cell neutralization were all AF-(16). Thus, the high titers of RhCMV-neutralizing antibodies in this non-transmitting cohort are expected. In contrast, the RhCMV-specific IgG infusions did not confer high titers of antibodies capable of mediating ADCP and ADCC functions, which may be a result of low levels of these types of antibodies in chronically infected animals from whom the IgG for these infusions was purified. Thus, while neutralizing responses were the only functional responses distinct between the CD4-depleted transmitting and non-transmitting dams, these findings may be a result of the experimental design and does not eliminate Fc mediated effector functions as potentially protective IgG responses.

A key caveat of this work, and most non-human primate studies, is limited group sizes. Thus, we included dams from prior studies of primary maternal infection in the rhesus macaque model to increase our sample size. This strategy has the advantage for this study of including animals with perturbations to their humoral responses, as CD4+ T cell depletion results in a delay in humoral responses and passive antibody infusion introduces a high titer of RhCMV-specific antibodies prior to infection (15, 16). These prior studies demonstrated that pre-existing antibodies can protect against vertical RhCMV transmission in the high risk setting of CD4-depletion (16). However, a caveat of including dams from multiple treatment groups is a potential confounding factor since all 3 of the dams receiving the high RhCMV-neutralizing IgG infusion were AF-negative and all 6 of the CD4-depleted dams without IgG infusion were AF-positive. Thus, this analysis is largely exploratory, and further experimentation of critical components of pre-existing immunity and cCMV transmission status would be necessary to confirm the trends observed.

While it is promising that we observed a similar rate of vertical CMV transmission in seronegative immunocompetent rhesus dams as has been observed in humans (3), the humoral and virological factors we investigated were largely not associated with AF status. Our findings underscore the need for further research to deconvolute the immune responses necessary for prevention of cCMV infection and ultimately to define targetable factors for vaccines or therapeutics. In the immunocompromised dams, we found that pre-existing antibodies can protect against vertical CMV transmission, suggesting that vaccination to elicit pre-existing IgG responses could be beneficial in prevention of cCMV in seronegative women. Innate immunity, T cell responses, and maternal-fetal interface immunity and inflammation are additional factors that may play influence vertical CMV transmission risk (30). Notably, CMV is also exceptionally adept at immune evasion as a large portion of its ∼235 kilobase genome is devoted to genes with immune evasion functions (31), so countering key immune evasins may additionally be a strategy for improving upon current preventive and therapeutic interventions. We have demonstrated that natural humoral immunity to primary infection during pregnancy will likely not be protective against congenital CMV infection, so developing effective strategies for establishing robust adaptive immunity prior to pregnancy is paramount to reducing the global burden of CMV-associated birth defects.

## Methods

### Animals

Indian-origin rhesus macaques were from a RhCMV-seronegative expanded specific pathogen–free (eSPF) colony housed at the Tulane Primate National Research Center or previously at the New England Primate Research Center and maintained in accordance with institutional and federal guidelines for the care and use of laboratory animals (32). We screened females for RhCMV-specific IgM and IgG via a whole virion ELISA to confirm that they were RhCMV-seronegative. These females, all between 4 and 9 years of age, were co-housed with RhCMV-seronegative males and screened for pregnancy every three weeks via abdominal ultrasound. Sonography was further used to approximate gestational date using the gestational sac size and crown-rump length of the fetus. A total of 27 RhCMV-seronegative dams were inoculated intravenously with RhCMV strains (1×10^6^ pfu for UCD52, UCD59 and FL RhCMV or 2-3×10^6^ TCID_50_ for 180.92) in late first/early second trimester (approximately 7–9 weeks gestation). One week prior to infection, 12 of the dams received 50 mg/kg recombinant rhesus CD4+ T cell–depleting antibody (CD4R1 clone; NIH Nonhuman Primate Reagent Resource) via intravenous infusion (15). Of the dams receiving the CD4-depletion antibody, 6 also received passive infusion of IgG purified from RhCMV seropositive animals. Three of these received a single infusion 1 hour before inoculation of 100mg/kg of IgG purified from animals screened for high RhCMV-binding via whole virion ELISA. The other three received a dose-optimized regimen of one infusion 1 hour prior to infection of 150 mg/kg and another at 3 days post infection of 100 mg/kg of IgG purified from animals screened for high RhCMV neutralization on epithelial cells (16). Blood was collected at least one timepoint before administration of any treatments, at the time of CD4+ T cell depletion, at the time of passive antibody infusions, and at the time of infection. The animals receiving passive antibody infusion were additionally sampled at 7–8 hours, 2 days and 4 days post infection. Dams in the new immunocompetent cohort were sampled at days 1 and 4 post infection in the first week. Otherwise, samples were collected on an approximately weekly basis in all groups. Amniocentesis was performed on a weekly to bi-weekly basis, when possible to perform safely by veterinary staff, until fetal loss (n = 5) or until hysterotomy in near-term of pregnancy (gestational weeks 19–22).

### Cell culture

Telomerized rhesus fibroblasts (TeloRFs) were maintained in Dulbecco’s modified Eagle medium (DMEM, Gibco) containing 10% heat-inactivated fetal bovine serum (FBS, Corning), 25 mM N-2-hydroxyethylpiperazine-N’-2-ethanesulfonic acid (HEPES, Gibco), 2 mM L-glutamine (Gibco), 50 U/mL penicillin and 50 μg/mL streptomycin (Gibco), 50 μg/mL gentamicin (Gibco), and 100 μg/mL geneticin (Gibco). Monkey kidney epithelial (MKE) cells were cultured in DMEM-F12 (Gibco) supplemented with 10% FBS, 25 mM HEPES, 2 mM L-glutamine, 1 mM sodium pyruvate (Gibco), 50 U/ml penicillin and 50 μg/ml streptomycin, and 50 μg/ml gentamicin. 293T cells used for transfection were maintained in DMEM supplemented with 10% FBS, 25 mM HEPES, 50 U/mL penicillin and 50 μg/mL streptomycin. THP-1 cells (ATCC) were maintained in RPMI 1640 medium (Gibco) supplemented with 10% FBS. NK92.rh.158I.Bb11 cells were cultured in Myelocult H5100 (STEMCELL) medium supplemented with 10mM HEPES, 1 mM sodium pyruvate, 50 U/ml penicillin and 50 μg/ml streptomycin, and 50 U/mL recombinant human IL-2 (PeproTech). Cell lines were generally passaged twice weekly and tested for mycoplasma contamination every 6 months.

### Virus growth

Stocks of epithelial-tropic UCD52 was produced by propagating from a seed stock on monkey kidney epithelial (MKE) cells for 2–4 passages and then once of telomerized rhesus fibroblasts (TeloRFs) to boost viral titer. Infection was performed in a low volume of reduced-FBS medium for at least 2 hours at 37°C and 5% CO_2_, a maintenance volume of media was added, and cells were incubated for up to 2 weeks. Virus was harvested when 90% cytopathic effect (CPE) was observed. Cells were collected by scraping and combined with the culture supernatant. Cells were pelleted by low-speed centrifugation and supernatant collected and placed on ice. Cell pellets were resuspended in infection media and combined, then subjected to either three freeze/thaw cycles or sonication. Large cell debris was pelleted by low-speed centrifugation. The supernatants were combined, passed through a 0.45-μm filter, overlaid onto a 20% sucrose cushion, and then ultracentrifuged at 70,000 g for 2 hours at 4°C using an SW28 Beckman Coulter rotor. Virus pellets were resuspended in DMEM containing 10% sucrose and titered on TeloRFs and/or MKE cells (depending on the intended use) using the TCID_50_ method.

FL-RhCMV was reconstituted in primary embryonal rhesus fibroblasts (1RFs) generated at the Oregon National Primate Research Center (ONPRC) by transfecting purified BAC DNA using Lipofectamine 3000 (Thermo Fisher Scientific). The cells were maintained in DMEM supplemented with 10% fetal bovine serum and antibiotics (1× Pen/Strep, Gibco), and grown at 37°C in humidified air with 5% CO_2_. 1RFs were passaged weekly until full CPE were observed. Cells and supernatants of one T-175 flask were harvested and used to infect 8 x T-175 flasks containing a confluent monolayer of 1RFs to produce a high titer stock. Cells and supernatants were harvested at maximal CPE and stored together over night at −80°C to release virus from the infected cells. The cellular supernatants were subsequently clarified by centrifugation at 7,500 × g for 15 min and the clarified culture medium was layered over a sorbitol cushion (20% d-sorbitol, 50 mM Tris [pH 7.4], 1 mM MgCl2). FL-RhCMV was pelleted by centrifugation at 22,000 rpm for 1 h at 4°C in a Beckman SW28 rotor and the resulting virus pellet was resuspended in DMEM, aliquoted and stored at −80°C until final use.

### Viral load

RhCMV DNA was quantified in maternal plasma and fetal amniotic fluid via qPCR. DNA was isolated from the amniotic fluid by using the QIAmp DNA minikit (Qiagen). We utilized a 45-cycle real-time qPCR reaction using 5′-GTTTAGGGAACCGCCATTCTG-3′ forward primer, 5′-GTATCCGCGTTCCAATGCA-3′ reverse primer, and 5′-FAM-TCCAGCCTCCATAGCCGGGAAGG-TAMRA-3′ probe directed against a 108bp region of the highly conserved RhCMV IE gene (19), using a standard IE plasmid to interpolate the number of viral DNA copies/mL of plasma or amniotic fluid. Vertical transmission was defined as 2 or more of 12 technical replicates above the limit of detection (1–10 copies per reaction) for each amniotic fluid sample. Two out of six replicates above the limit of detection was required to report a positive result for each plasma sample.

### Serology assays

*RhCMV-specific IgG antibody kinetics* were measured in plasma by enzyme-linked immunosorbent assays (ELISAs) using either whole-virion preparations of RhCMV strains UCD52 or purified glycoprotein preparations of gB or pentameric complex as described previously (16). Briefly, high-binding 384-well ELISA plates (Corning) were coated with 15 μL/well of antigen overnight at 4°C. For whole virion ELISAs, filtered RhCMV UCD52 was diluted to 5,120 PFU/mL in phosphate buffered saline with Mg^2+^ and Ca^2+^ (PBS+) for coating, while soluble glycoprotein preparations were coated at 2 μg/mL in 0.1M sodium bicarbonate. Following overnight incubation, plates were blocked for 2 hours with blocking solution (PBS+, 4% whey protein, 15% goat serum, 0.5% Tween-20), and then 3-fold serial dilutions of plasma (1:30 to 1:65,610) were added to the wells in duplicate for 1.5 hours. Plates were then washed using an automated plate washer (BioTek) and incubated for 1 hour with mouse anti-monkey IgG HRP-conjugated secondary antibody (Southern Biotech, clone SB108a) at a 1:5,000 dilution or anti-monkey IgM HRP-conjugated secondary antibody (Rockland) at 1:8,000. After two washes, SureBlue Reserve TMB Microwell Peroxidase Substrate (KPL) was added to the wells for 7 minutes for whole virion ELISAs and 3.5 minutes for glycoprotein ELISAs, and the reaction was stopped by addition of 1% HCl solution (KPL). Plates were read at 450 nm and reported as ED_50_ or area-under-the-curve of the dilution series. The lower threshold for antibody reactivity was considered to be 3 SDs above the average 450 nm optical density measured for RhCMV-seronegative samples at the starting plasma dilution (1:30). ED_50_ was calculated as the sample dilution where 50% binding occurs by interpolation of the sigmoidal binding curve using four-parameter logistic regression, and this value was set to half of the starting dilution (15) when the 1:30 dilution point was negative by our cutoff and when the calculated ED_50_ was below 15.

*Neutralization of RhCMV on fibroblasts and epithelial cells* was monitored as previously described (16). Briefly, serial dilutions (1:10 to 1:30,000 or 1:30 to 1:65,610) of heat-inactivated rhesus plasma were incubated with FL RhCMV (13) or RhCMV UCD52 (MOI = 1) for 1–2 hours at 37°C. The virus/plasma dilutions were then added in duplicate to 384-well plates containing confluent cultures of TeloRF or MKE cells, respectively, and incubated at 37°C for 24 or 48 hours, respectively. Infected cells were then fixed for 20 minutes in 10% formalin and processed for immunofluorescence with 1 μg/mL mouse anti-RhCMV pp65B or 5 ug/mL mouse anti-Rh152/151 monoclonal antibody followed by a 1:500 dilution of goat anti-mouse IgG-Alexa Fluor 488 antibody (Invitrogen). Nuclei were stained with DAPI for 10 minutes (Invitrogen). Infection was quantified in each well by automated cell counting software using one of the following instruments at 10X magnification: (a) a Nikon Eclipse TE2000-E fluorescence microscope equipped with a CoolSNAP HQ-2 camera, (b) a Cellomics CX5 or Arrayscan VTI HCS instrument, or (c) a Molecular Devices ImageXpress Pico. Subsequently, the ID_50_ was calculated as the sample dilution that caused a 50% reduction in the number of infected cells compared with wells treated with virus only. If the ID_50_ was below the limit of detection (e.g., every value in the series resulted in a high level of infection above 50%), we set the ID_50_ to half of the first dilution. Similarly, for samples above the upper limit of detection (e.g., all points in the series resulted in levels of infection below 50%), we set the ID_50_ at the value of the last dilution point.

*Cell-associated gB binding* was measured using a gB-transfected-cell binding assay (29). 293T cells at 50– 80% confluence were co-transfected with 1 μg of each GFP (gift of M. Blasi, Duke University) and UCD52 gB (Ampicillin-resistant pcDNA3.1+ vector) using the Effectene transfection kit (Qiagen) and incubated for 48 hours. Following transfection, GFP expression was confirmed using a fluorescent microscope prior to harvest. Cells were then dissociated from culture flasks using TrypLE (Gibco), and 100,000 cells per well were added to 96-well V-bottom plates (Corning). Transfected cells were incubated with plasma samples at a 1:600 dilution in duplicate for 2 h. Cells were then washed and resuspended in far red live/dead stain (Invitrogen) at 1:1000 and incubated for 20 minutes. Following another wash, cells were resuspended in 1:200 of anti-monkey IgG-PE (Southern Biotech) for 25 minutes at RT. Cells were then washed twice and fixed for 20 minutes in 10% formalin and resuspended in PBS. Fluorescence was measured using a BD LSRII or Fortessa flow cytometer, and data is reported as the percent of the live population that is GFP and PE double positive.

*Antibody-dependent cellular phagocytosis* was assessed by conjugation of concentrated RhCMV UCD52 to Alexa Fluor 647 using NHS-ester reaction (Invitrogen), which was allowed to proceed in the dark with constant agitation for 1–2 hours. The reaction was quenched by the addition of pH 8.0 Tris-HCl, and the conjugated virus was diluted to 7×10^6^ PFU/mL. Equal amounts (10 μL) of virus and plasma samples in duplicate at a 1:30 dilution were combined in a 96-well U-bottom plate (Corning) and incubated at 37°C for 2 hours. THP-1 monocytes were then added at 50,000 cells per well. Plates were spun for 1 hour at 1200xg at 4°C in a spinoculation step and then transferred to a 37°C incubator for an additional 1 hour. Cells were then washed and stained with aqua live/dead (Invitrogen) at 1:1000 for 20 minutes. Following another wash step, cells were fixed for 20 minutes in 10% formalin and resuspended in PBS. Fluorescence was measured using a BD LSRII or Fortessa flow cytometer, and data is reported as the percent of the live population that was AF647+.

*Antibody-dependent cellular cytotoxicity* was measured by NK cell degranulation using NK92.rh.158I.Bb11, an engineered NK cell line expressing the Ile^158^ variant of rhesus macaque CD16 (33). Confluent TeloRF cells were infected with 1 MOI UCD52 RhCMV in low serum (5%) medium in T75 flasks (ThermoFisher). Mock infection was performed in parallel. After 24 hours of infection, cells were dissociated from the flask using TrypLE (Gibco), and 50,000 target cells were seeded in each well of a 96-well flat bottom tissue culture plate (Corning). Cells were incubated for 16–20 hours to allow them to adhere. Then, the target cell medium was removed, and to each well, 50,000 NK92.Rh.CD16-Bb11 cells were added with plasma samples at 1:25 dilution or purified IgG from either seropositive or seronegative plasma donors at 50–250 µg/mL in R10 (RPMI1640 containing 10% FBS), plated in duplicate. The same samples were added to mock-infected target cells for background subtraction. Brefeldin A (GolgiPlug, 1:1000, BD Biosciences), monensin (GolgiStop, 1:1500, BD Biosciences), and CD107a-FITC (BD Biosciences, 1:40, clone H4A3) were added to each well for a total of 100μL and co-incubated for 6 hours at 37°C and 5% CO_2_. NK cells were then collected and transferred to a 96-well V-bottom plate (Corning). Cells were pelleted, washed with PBS, and then resuspended in aqua live/dead diluted 1:1000 for a 20-minute incubation at RT. Cells were washed with PBS + 1% FBS and stained with CD56-PECy7 (BD Biosciences, clone NCAM16.2) and CD16 PacBlue (BD Biosciences, clone 3G8) for a 20-minute incubation at RT. Cells were washed twice then resuspended in 10% formalin fixative for 20 minutes or 1% formalin overnight. Cells were then resuspended in PBS for acquisition. Events were acquired on a BD Symphony A5 using the high-throughput sampler (HTS) attachment, and data analysis was performed using FlowJo software (v10.8.1). Data is reported as the background-subtracted % of CD107a+ live NK cells (singlets, lymphocytes, aqua blue–, CD56+, CD107a+) for each sample.

### Statistical analysis

For each biomarker we calculated the area under the curve from day 0 to day 21. In instances where there was not a day 0 value, we either A) carried forward the last value before infection (if existed) or B) linearly back-extrapolated from the first two values after infection. When there were no day 21 values, we either A) linearly extrapolated between last value before day 21 and first value after day 21 (if existed) or B) carried forward last value before day 21 (if no value post day 21). We performed exact Wilcoxon-rank sum tests using the coin package in R (34). Multiple testing correction was done using Benjamini-Hochberg FDR method with significance deemed as an adjusted p-value less than 0.2 and an unadjusted less than 0.05 (35). In the combined analyses, we used the coin package again but performed the analysis stratified by immune status (this assumes similar association across groups). PCA was calculated in R using prcomp.

## Acknowledgements

The authors would first like to acknowledge the rhesus macaques that were used in this study and thank the faculty and staff of the Departments of Veterinary Medicine and Collaborative Research at the Tulane National Primate Research Center (TNPRC) for their excellent care of our research animals. We would like to thank former lab staff, namely Kristy Bialas and Eduardo Cisneros de la Rosa, for the contribution of previously published ELISA and neutralization data included in this analysis. We acknowledge the Molecular Virology Core at Oregon National Primate Research Center (ONPRC) for performing viral productions, supported by NIH grant award P51OD011092. We are additionally grateful to the NIH Nonhuman Primate Reagent Resource (R24 OD010976, and NIAID contract HHSN 272201300031C) which provided the CD4-depleting antibody used in the historical studies. We wish to acknowledge support from the Biostatistics, Epidemiology and Research Design (BERD) Methods Core funded through Grant Award Number UL1TR002553 from the National Center for Advancing Translational Sciences (NCATS), a component of the NIH. Research funds were provided by NIH/NIAID 1P01AI129859-01A1 and NIH/NICHD Director’s New Innovator grant to S.R.P. (DP2HD075699), and the TNPRC base grant NIH/NCRR (OD011104). C.E.O. was supported by T32-CA009111 and 3P01AI129859-02S1. The funders had no role in study design, data collection and interpretation, decision to publish, or the preparation of this manuscript. The content is solely the responsibility of the authors and does not necessarily represent the official views of the NIH.

